# Structural Basis for Accommodation of Emerging B.1.351 and B.1.1.7 Variants by Two Potent SARS-CoV-2 Neutralizing Antibodies

**DOI:** 10.1101/2021.02.21.432168

**Authors:** Gabriele Cerutti, Micah Rapp, Yicheng Guo, Fabiana Bahna, Jude Bimela, Eswar R. Reddem, Jian Yu, Pengfei Wang, Lihong Liu, Yaoxing Huang, David D. Ho, Peter D. Kwong, Zizhang Sheng, Lawrence Shapiro

## Abstract

Emerging SARS-CoV-2 strains, B.1.1.7 and B.1.351, from the UK and South Africa, respectively show decreased neutralization by monoclonal antibodies and convalescent or vaccinee sera raised against the original wild-type virus, and are thus of clinical concern. However, the neutralization potency of two antibodies, 1-57 and 2-7, which target the receptor-binding domain (RBD) of spike, was unaffected by these emerging strains. Here, we report cryo-EM structures of 1-57 and 2-7 in complex with spike, revealing each of these antibodies to utilize a distinct mechanism to bypass or accommodate RBD mutations. Notably, each antibody represented a response with recognition distinct from those of frequent antibody classes. Moreover, many epitope residues recognized by 1-57 and 2-7 were outside hotspots of evolutionary pressure for both ACE2 binding and neutralizing antibody escape. We suggest the therapeutic use of antibodies like 1-57 and 2-7, which target less prevalent epitopes, could ameliorate issues of monoclonal antibody escape.

## Introduction

Severe acute respiratory syndrome coronavirus 2 (SARS-CoV-2), the causative agent for Coronavirus Disease 2019 (COVID-19), emerged in 2019, rapidly establishing an ongoing worldwide pandemic with over 100 million infected and over two million dead at the current time (Callaway et al., 2020; Cucinotta and Vanelli, 2020; Dong et al., 2020). The possible emergence of mutant strains of SARS-CoV-2 with improved transmissibility, virulence, or ability to evade human immunity has been a major concern (Baric, 2020). The SARS-CoV-2 lineage known as B.1.1.7 emerged in September 2020 in South East England, quickly becoming the dominant variant in the UK, and subsequently spreading to over 50 countries, potentially due to enhanced transmissibility (Andrew Rambaut et al., 2020). The B.1.1.7 strain contains the early spike mutation D614G, now common to most SARS-CoV-2 lineages, as well as eight additional spike mutations including two deletions (69-70del & 144del) in the N-terminal domain (NTD), a single mutation (N501Y) in receptor-binding domain (RBD) and (A570D) in SD1, two mutations (P681H and T716I) near the furin cleavage site, and two mutations in S2 (S982A and D1118H). Another emerging lineage, SARS-CoV-2 B.1.351, appeared late in 2020 in Eastern Cape, South Africa (SA) (Tegally et al., 2020), and also become dominant locally, again raising the possibility of increased transmissibility. B.1.351 contains 9 spike mutations in addition to D614G, including a cluster of mutations (e.g., 242-244del & R246I) in NTD, one mutation (A701V) near the furin cleavage site, and three mutations (K417N, E484K, & N501Y) in RBD.

Recent studies have shown that some of these new variants impede the function of some SARS-CoV-2 neutralizing antibodies, with most NTD-directed neutralizing antibodies showing a near complete loss of potency against either the B.1.351 or B.1.1.7 strains (Wang et al., 2021; Wibmer et al., 2021). With respect to RBD-directed antibodies, however, while the most prevalent classes of multi-donor RBD-directed antibodies, originating from the VH3-53/66 and VH1-2 genes (Banach et al., 2021; Rapp et al., 2021; Yuan et al., 2020) are generally inhibited by mutations in RBD (K417N and N501Y for VH3-53/66 class antibodies and E484K for VH1-2 class antibodies) (Wang et al., 2021). Fortunately, many RBD-directed antibodies not from these frequent classes retain their activity.

In a companion manuscript (Wang et al., 2021), we screened monoclonal antibodies and found two super potent RBD-directed antibodies, 1-57 and 2-7 (0.008 and 0.003 μg/mL of live virus neutralization IC50 for 1-57 and 2-7, respectively), whose neutralization was unaffected by the emerging B.1.351 and B.1.1.7 strains. To understand how these antibodies accommodate the mutations present in B.1.351 and B.1.1.7 strains, we determined their cryo-EM structures in complex with spike. We used the 52 currently known complex structures of RBD-directed neutralizing antibody to create a residue-level map of antibody recognition frequency. This analysis suggested mutations in RBD to emerge under evolutionary pressure from both antibody-escape and binding to the ACE2 receptor on host cells. We suggest that neutralizing antibodies such as 1-57 and 2-7 that target epitopes with few selection hotspots may result in less viral escape and lead to more effective therapeutic strategies.

## Results

### Antibody 1-57 utilizes a hydrophilic pocket to accommodate mutation E484K in emerging strains

From complexes of SARS-CoV-2 spike -stabilized by 2P mutations (Wrapp et al., 2020) -with the antigen-binding fragment (Fab) of antibody 1-57, we collected single particle data on a Titan Krios microscope, yielding a cryo-EM reconstruction to an overall resolution of 3.42 Å (**Figure 1A, Figures S1 and S2, Table S1**). A single conformational state with three Fabs per trimer, each bound to an RBD in the ‘down’ conformation, was identified. Recognition of RBD by 1-57 was dominated by the heavy chain, which buried 533.7 Å^2^ surface area, with a smaller 223.3 Å^2^ contribution by the light chain. The 21 amino acid-long CDR H3 formed a β-hairpin stabilized by a disulfide bond between Cys100a and Cys100f, provided the primary interactions with RBD, with additional contributions from CDR L1 and L2 (**Figure 1C, left panel**). CDR H3 formed mainly hydrophobic contacts with RBD residues Tyr351, Leu452, Phe490 and Leu492. The light chain contributed to RBD recognition mainly through hydrogen bonds formed between the hydroxyl groups of Ser30, Tyr32 and Ser53 in CDR L1 and L2 and residues Gln493, Ser494 and Gln498 in RBD respectively (**Figure 1C, right panel**). Antibody 1-57 also displayed a small quaternary interaction, as Asn100c on the tip of CDR H3 hydrogen bonds with both Thr470 in a neighboring RBD and glycan *N*165 in a neighboring NTD **(Figure 1C, middle panel**).

**Figure 1.**
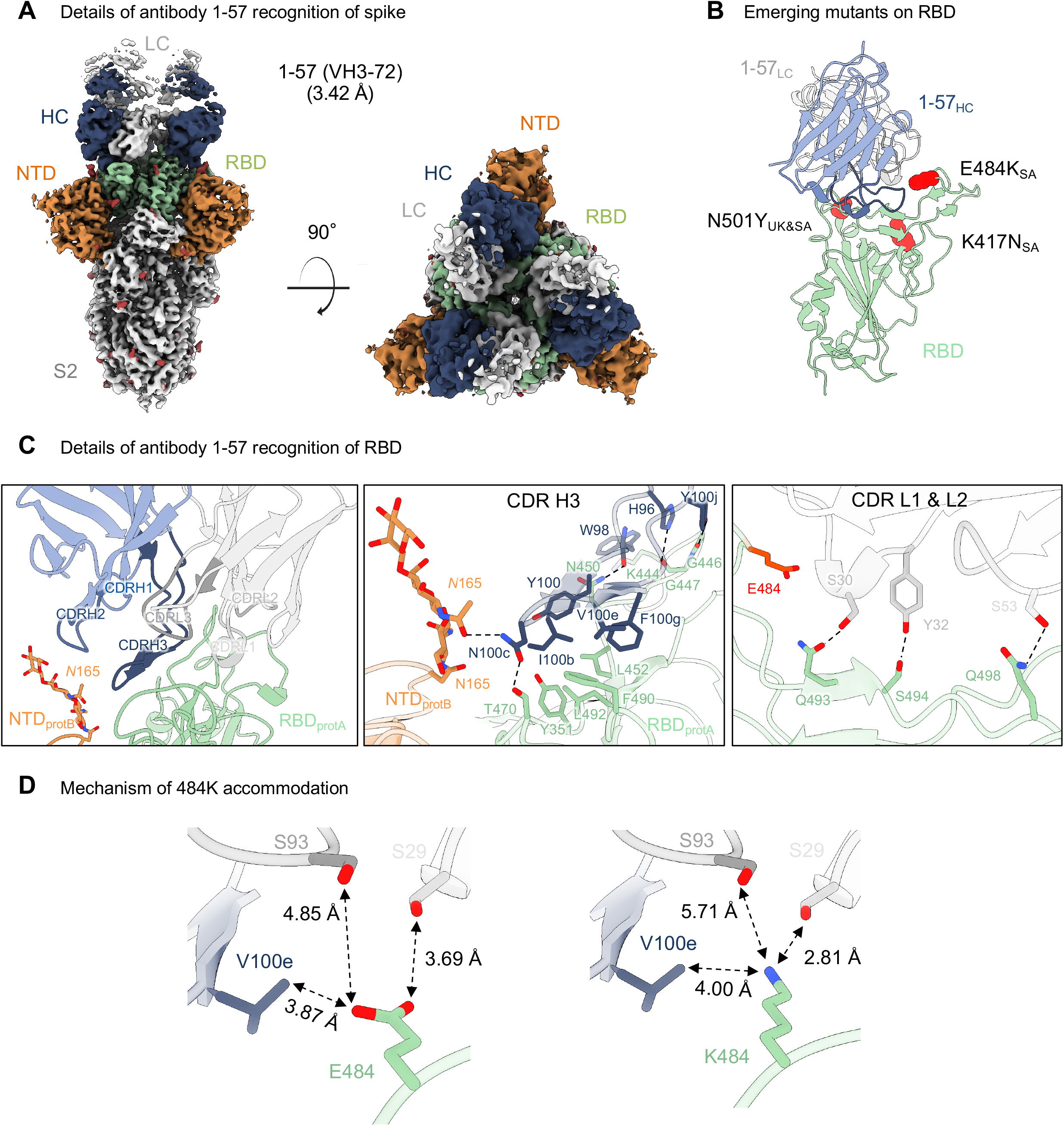
Antibody 1-57 utilizes a hydrophilic pocket to accommodate mutation E484K in emerging strains. (A) Cryo-EM reconstruction for spike complex with antibody 1-57 from two orthogonal views; a single conformation with all RBDs down is observed. NTD is shown in orange, RBD in green, glycans in red, antibody heavy chain in blue and light chain in gray. (B) Domain level view of 1-57 in complex with RBD, with the emerging mutants highlighted in red. (C) Details of antibody 1-57 recognition of RBD showing the overall interface (left panel), recognition by CDR H3 (middle panel) and recognition by CDR L1 and L2 (right panel). CDR H1, H2, H3 are colored in shades of blue; CDR L1, L2, and L3 are colored in shades of gray. E484 is highlighted in bright red (right panel). Nitrogen atoms are colored in blue, oxygen atoms in red; hydrogen bonds (distance <3.2 Å) are represented as dashed lines. (D) Expanded view of the E484 environment at the interface with 1-57 (left panel) and modeling of K484 (right panel) suggest a mechanism of antibody 1-57 accommodation of the E484K mutation; colored as in (B). See also Table S1, Figure S2.

With respect to the three RBD mutations in the UK and South Africa strains, only residue E484K of the South African strain was near the binding site of 1-57 (**Figure 1B**). Despite its proximity to the epitope, however, Glu484 did not interact significantly with 1-57, as the amino acids in the immediate vicinity – Val100e from the heavy chain, and Ser29 and Ser93 from the light chain – were too distant (**Figure 1D, left panel**). Structural modeling of the E484K mutation showed that a Lys residue was geometrically compatible with 1-57 binding; modelling of K484 with a high frequency rotamer showed that the distance between the amino group of K484 and the side chain of Ser29 was compatible with a hydrogen bond (2.81 Å) (**Figure 1D, right panel**).

### Structural basis of antibody 2-7 accommodation of mutation N501Y in emerging strains

Cryo-EM analysis of the Fab from antibody 2-7 in complex with SARS-CoV-2 spike produced a reconstruction with 3 Fabs bound to a single spike, and was refined to an overall resolution of 3.72 Å (**Figure 2A, Figures S1 and S3, Table S1**). Only one conformation was observed, with Fabs bound to two RBDs in the ‘up’ conformation and one in the ‘down’ conformation. Due to extensive conformational heterogeneity, the ‘up’ RBDs could not be resolved to high resolution, thus the ‘down’ RBD was the focus of structural analysis.

**Figure 2.**
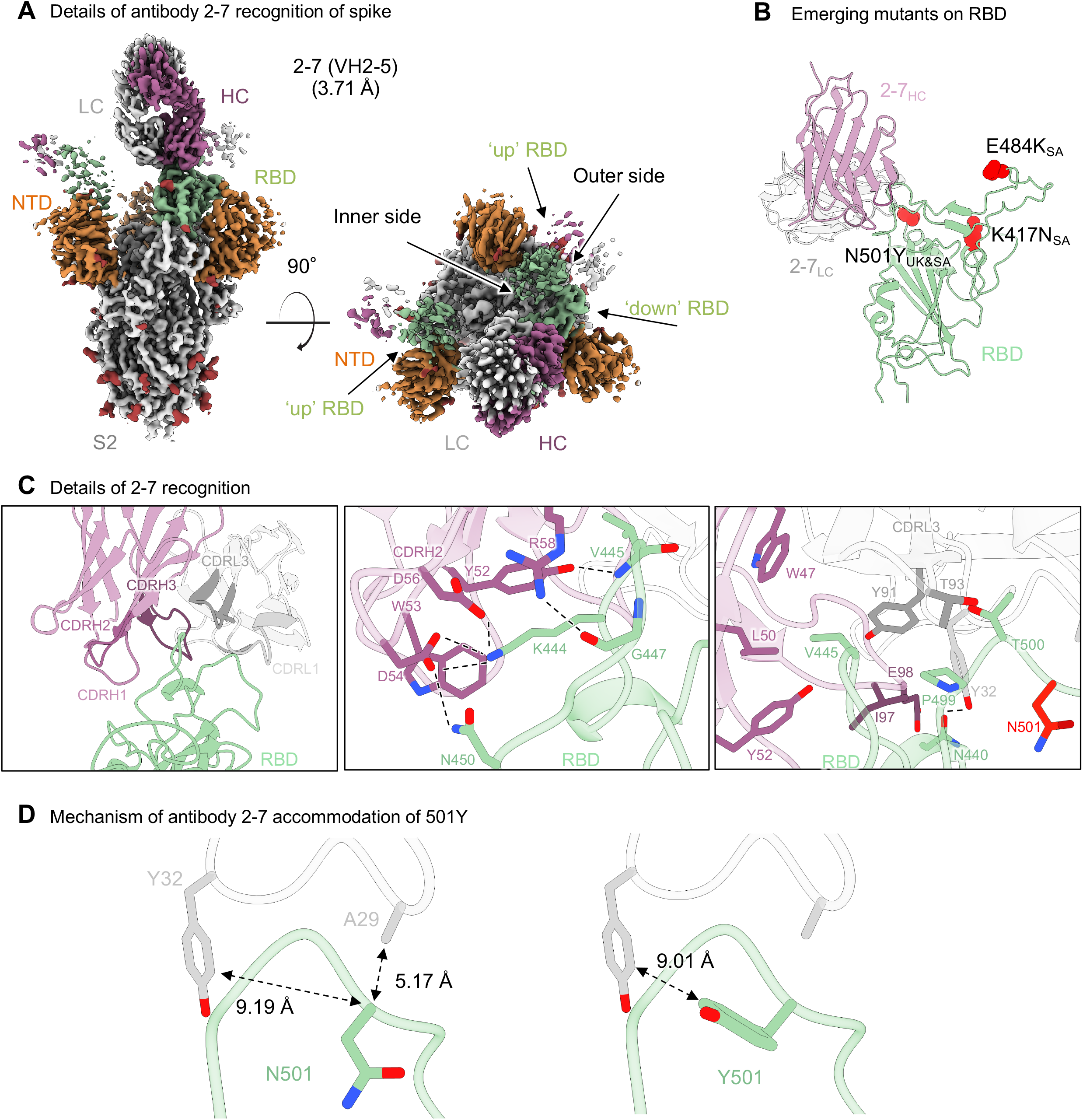
Structural basis of antibody 2-7 accommodation of mutation N501Y in emerging strains. (A) MCryo-EM reconstruction for spike complex with antibody 2-7 from two orthogonal views; a single conformation with 1 RBD down and two RBDs up is observed. NTD is shown in orange, RBD in green, glycans in red, antibody heavy chain in magenta and light chain in gray. (B) Domain level view of 2-7 in complex with RBD, with the emerging mutants highlighted in red. (C) Details of antibody 2-7 recognition of RBD showing the overall interface (left panel), recognition by CDR H2 (middle panel) and recognition by CDR L1 and L3 (right panel). CDR H1, H2, H3 are colored in shades of magenta; CDR L1, L2, and L3 are colored in shades of gray. N501 is highlighted in bright red (right panel). (D) Expanded view of the N501 environment at the interface with 2-7 (left panel) and modeling of Y501 (right panel) suggest a mechanism of antibody 2-7 accommodation of the N501Y mutation; colored as in (B). See also Table S1, Figure S3.

Recognition of RBD by antibody 2-7 was dominated by interactions proximal to the RBD loops formed by residues 438-451 and 495-502 (**Figure 2C, left panel**). CDR H2 formed an extensive network of hydrogen bonds and salt bridges (**Figure 2C, middle panel**). RBD residue Lys444 formed salt bridges with Fab heavy chain residues Asp56 and Asp54. Additionally, CDR H2 residues Asp54, Tyr52, and Arg58 formed hydrogen bonds with RBD residues Asn450, the backbone amine of Val445, and the backbone carbonyl of Gly447, respectively. Val445 at the apex of loop 438-451 was at the center of a hydrophobic pocket that included Pro499 on RBD that accommodated several residues on the heavy chain (Trp47, Leu50, Tyr52, and Ile97) and one on the light chain (Tyr91). Light chain residue Tyr32 also formed a hydrogen bond with the RBD via residue N440 (**Figure 2C, right panel**).

With respect to the three mutated positions in the UK and South Africa variants, antibody 2-7 bound near only N501, but the sidechain of N501 pointed away from the antibody (**Figure 2B and Figure 2D**). While some conformational change of the 495-502 loop would be expected in the context of the N501Y mutation, this loop contributed only 225 Å^2^ out of 736 Å^2^ and contained few residues that form significant interactions with the Fab.

### Genetic and epitope similarities to other SARS-CoV-2 neutralizing antibodies

The heavy chain of 1-57 derived from the VH3-72*01 gene with a CDR H3 of 21amino acids, and the light chain was from KV3-20*01, with a 9 residue CDR L3 (**Figure S3**). Antibody 2-7 utilized VH2-5*02 with an 11 amino acid CDR H3, and LV2-14*01 with a 9 residue CDR L3. Both antibodies thus utilized heavy chain genes with relatively low frequencies in SARS-CoV-2 specific neutralizing antibody repertoire (Rapp et al., 2021), suggesting that antibodies with genetic features similar to 1-57 and 2-7 may not appear frequently in human response to SARS-CoV-2.

To understand whether the binding orientations of antibodies 2-7 and 1-57 were common among RBD-directed antibodies, we superposed structures of RBDs from the 52 RBD-directed antibody complex structures deposited in the PDB and measured root mean square deviations (RMSD) between antibodies. Overall, clustering of the 52 antibodies using pairwise RMSD showed that antibodies 1-57 and 2-7 did not cluster with the most frequent VH3-53 and VH1-2 antibody classes (**Figure 3A**). Antibody 2-7 was grouped with REGN10987, which was a component of a SARS-CoV-2 therapeutic cocktail (Hansen et al., 2020). The binding orientation of antibody 1-57 was similar to antibodies P2B-2F6 and CV07-270. However, these antibodies utilize paratopes different from 2-7 and 1-57 for RBD recognition, probably because each antibody has a unique genetic origin.

**Figure 3.**
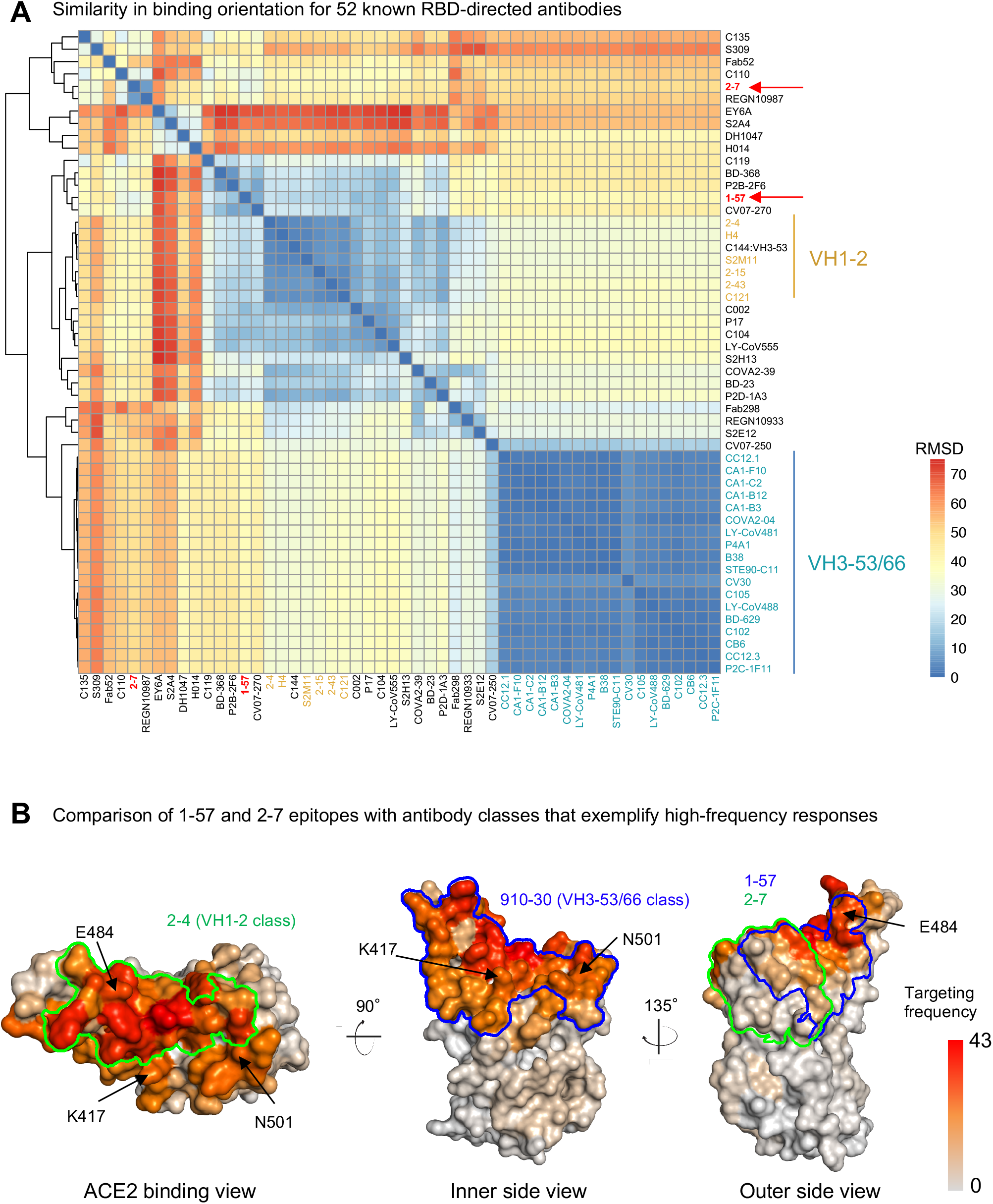
Antibodies 1-57 and 2-7 exemplify rare responses, suggesting that mutations against these antibodies have low selection pressure. (A) Analysis of 52 known RBD-directed neutralizing antibodies indicates that 1-57 and 2-7 approach RBD with angles distinct from prevalent antibody classes. (B) Per residue frequency recognized by 52 known RBD-directed antibodies. VH1-2 and VH3-53/66 antibody classes recognize RBD residues with high targeting frequency. 1-57 and 2-7 recognize RBD residues with low targeting frequencies.

During *in vivo* viral infection, epitopes frequently targeted by antibody may impose strong selection pressure for viral escape. We used the 52 neutralizing antibodies in the PDB to estimate the frequencies of the epitopes of 1-57 and 2-7 being recognized by assuming that these antibodies recapitulate *in vivo* recognition frequencies of RBD epitopes. Briefly, for each antibody, we identified epitope residues and calculated the frequency of each RBD residue being recognized by antibody. The analysis revealed that the epitope residues of 1-57 and 2-7 showed lower antibody recognition frequencies (about 11.2 and 19.8 antibodies per residue in average for 2-7 and 1-57, respectively) compared to those targeted by the prevalent antibody classes (about 27.4 and 25.9 antibodies per residue in average for VH1-2 and VH3-53 class, respectively, **Figure 3B**), suggesting that 1-57 and 2-7 epitopes are relatively less targeted antigenic sites.

### Antibody escape mutations may result from ACE2-binding and antibody-escape selection pressure

Analysis of RBD mutations observed in circulating SARS-CoV-2 sequences revealed many emerging mutations (**Figure 4A**). To understand whether the emerging mutations affect antibodies 1-57 and 2-7 recognition, we analyzed structural locations of the top 10 most frequent RBD mutations. We found that while antibodies 1-57 and 2-7 are not affected by the three mutations observed in the UK and SA strains, several RBD mutations may affect antibodies 1-57 and 2-7. For example, mutations at position 452 and 494 were within the antibody 1-57 binding footprint (**Figure 4B**). Mutations at position 439 may affect 2-7 binding. However, further experiments will be required to assess such effects.

**Figure 4.**
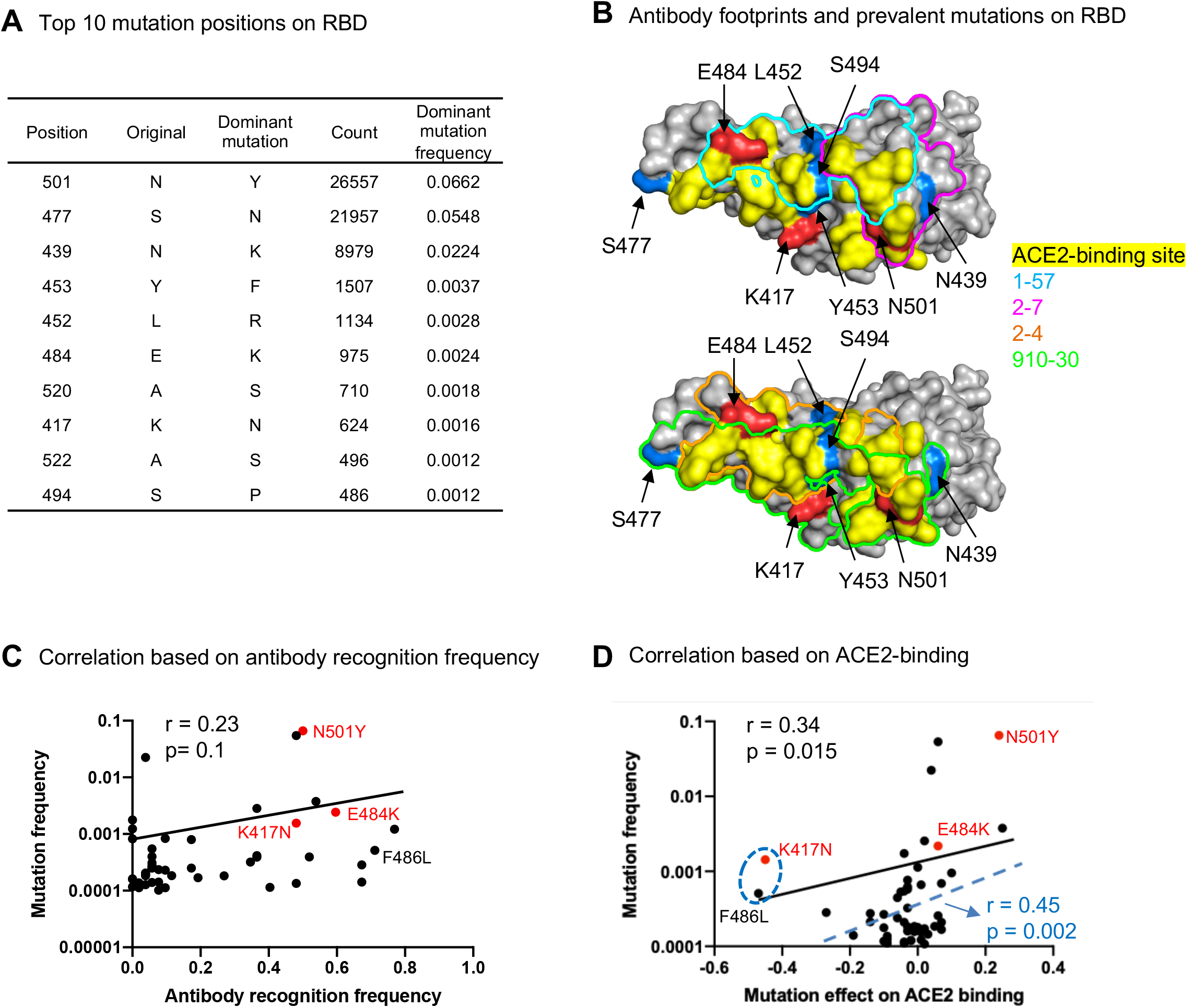
Prevalent emerging mutations appear to arise at epitopes of prevalent neutralizing response, suggesting that resistance mutants might arise less frequently to rare responses such as antibodies 1-57 and 2-7. (A) Most frequent mutations and positions observed in circulating SARS-CoV-2 strains. (B) Location of prevalent RBD mutations and antibody footprints. (C) Correlation between per residue antibody recognition frequency and top 50 RBD position mutations (D) Correlation between mutation effect on ACE2 binding and its frequency. The blue dashed line, r value and p value represent the fit after removing the 2 ‘outliers’, K417N and F486L mutations.

Studies have proposed that many SARS-CoV-2 escape mutations are selected by neutralizing antibodies (Baum et al., 2020; Ku et al., 2021). However, it is unclear whether the observed antibody resistant mutations are all the results of antibody selection. To shed light on this, we calculated correlation between positional mutation frequency and antibody recognition frequency by assuming that the estimated per RBD residue targeting frequency reflects *in vivo* frequency of antibody recognition and that the most frequently targeted RBD residues may undergo strong selection pressure to mutate. To our surprise, we only observed a very weak correlation (**Figure 4C**, r = 0.23, p = 0.1) albeit some of the most frequently targeted residues do have higher mutation frequencies. Because RBD is always under selection pressure from ACE2 binding for entry (Starr et al., 2020), we next examined whether the emerging RBD mutations were selected by enhancing ACE2 binding. We calculated the correlation between frequencies of RBD mutations and their normalized effects on ACE2 binding affinity obtained from deep mutational scanning data (Starr et al., 2020), which showed a significant correlation (**Figure 4D**, Pearson’s r = 0.34, p = 0.015). The correlation coefficient could be further increased significantly when removing two ‘outlier’ mutations, K417N and F486L, both of which reduce ACE2 binding affinity significantly (**Figure 4D**). Further analysis of the antibody escape mutations showed that the frequent antibody escape mutations we observed may arise from selection advantage for both ACE2 binding and antibody escape. For example, the most frequent N501Y could improve the binding affinity of ACE2 as well as impair recognition of the most frequent VH3-53 class antibodies (Wang et al., 2021). The E484K mutation – detected in B.1.351 and recently emerging strains P.1 and P.2 (Nuno R. Faria et al., 2021) – impairs recognition of numerous antibodies and also weakly improves ACE2 binding affinity. However, the frequent mutation K417N, resulted in escape of the prevalent VH3-53 derived antibody class (Wang et al., 2021), and may be selected predominantly by antibody pressure. Despite the ability of 5-7 and 2-17 to still potently neutralize emerging strains that carry N501Y, K417N and E484K mutation, we observed that individual mutations at 452, 494, and 439 (**Figure 4B**)– may affect 1-57 and 2-7 recognition -do not or only weakly enhance ACE2 binding (Starr et al., 2020). Thus, these mutations may not be selected by ACE2 binding enhancement, but further experiments are required to understand whether combinations of these individual mutations could have beneficial advantage.

Our analysis suggests that potent antibodies 2-7 and 1-57 may not put much selection pressure on circulating strains, thus resistance mutants to these two antibodies are less likely to arise. Overall, due to the diversity of emerging strains, different antibody cocktails become more and more important, and 2-7 and 1-57 are decent candidates for therapeutic development.

## Discussion

The emergence of mutant SARS-CoV-2 strains and their impact on the function of neutralizing antibodies has become a major concern. The B.1.1.7 (UK) and B.1.351 (South Africa) variants contain mutations that evade the most frequently elicited classes of RBD-directed neutralizing antibodies from the VH1-2 (B.1.351) and VH3-53/66 classes (B.1.1.7) (Ku et al., 2021; Wang et al., 2021). Both antibody classes recognize predominantly the RBM, which is mutated convergently in many emerging viral lineages (e.g. K417T, E484K, and N501Y in both B.1.351 and B.1.1.28 (Brazil)). In this study, we report the structures and accommodation mechanisms for two potent SARS-CoV-2 neutralizing antibodies, 1-57 and 2-7, which are not impaired by the mutations in the B.1.1.7 and B.1.351 lineages.

Quantifying epitope-targeting frequency is important for understanding mechanisms of antibody induced viral escape. Despite an algorithm having been developed to estimate linear epitope-targeting frequency by serum antibodies in a high-throughput way (Shrock et al., 2020), it is still difficult to measure the epitope-targeting frequency of antibodies recognizing three dimensional epitopes. Methods such as competition ELISA can only reveal frequencies of targeted epitope regions with no residue-level information. Here we use a method to roughly estimate residue-level frequencies for antibody recognition of RBD based on the 52 currently known atomic-level structures of neutralizing antibody-RBD complexes. These antibodies, identified in separate studies of infected donors from around the world (Barnes et al., 2020; Hansen et al., 2020; Lv et al., 2020; Pinto et al., 2020) were generally chosen for study due to their neutralization potency. Overall, these studies identified antibodies mainly targeting RBD (Brouwer et al., 2020; Shi et al., 2020; Tortorici et al., 2020; Wu et al., 2020), with some NTD-directed neutralizers (Cerutti et al., 2021; Chi et al., 2020), with antibodies of the same class often identified in multiple studies. The residue-level antibody-interaction frequencies we present for RBD are likely to represent a reasonable estimate of relative frequencies, but only for the most potent and most frequent classes. Thus, the calculated residue-level antibody-interaction frequency may not reflect epitope-targeting frequencies of weakly-or non-neutralizing antibodies, which are a significant portion of antibody response to SARS-CoV-2 (Liu et al., 2020; Robbiani et al., 2020). It is also unclear whether weakly-or non-neutralizing antibodies impose selection pressure on RBD epitopes. Therefore, the weak correlation between antibody-interaction frequency and mutation frequency could be due to multiple confounding factors not included. Nonetheless, many of the identified antibody-interaction hotspots are within or close to the ACE2 binding site. Linear-peptide epitopes around these hotspots are also recognized by numerous serum antibodies (Shrock et al., 2020), confirming that the hotspots we identified are frequently targeted.

The study of mechanisms of mutant accommodation by antibodies 1-57 and 2-7 shed insights on effective therapeutic strategies. Studies have shown that persistent SARS-CoV-2 infection can last for months in immunocompromised human individuals (Avanzato et al., 2020; Choi et al., 2020). During such a long acute infection period, the arm race between the immune system and virus results in many viral mutations to be accumulated in spike to escape antibody neutralization. Because the human neutralizing antibody response frequently and convergently targets the ACE2 receptor-binding region on RBD, there is strong positive selection pressure to mutate this region. In the meantime, our results showed that the most frequent mutations tend to enhance ACE2 binding, suggesting additional selection pressure underlying SARS-CoV-2 evolution. Because most RBD mutations are detrimental to ACE2 binding (Starr et al., 2020), the virus may have a limited mutation space for escaping antibody neutralization. This may explain the rapid spread of convergent RBD mutations in different emerging SARS-CoV-2 lineages (e.g. E484K mutation in Brazilian P.1, SA B.1.351, and US B.1.526 lineages) (Sabino et al., 2021; West et al., 2021). For therapeutics, antibodies that can either tolerate mutations in the ACE2-binding region or recognize epitopes outside this region are critical for protection. Here, 1-57 and 2-7 bind epitopes that are not predominantly focused on the ACE2-binding region and also accommodate RBD mutations. A residue-level interaction-frequency analysis of RBD interaction with all currently known RBD-directed neutralizing antibodies of structure revealed them to represent relatively low frequency antibody response, which may not form strong selection pressure on the epitope regions. Therapeutics developed from such neutralizing antibodies could lead to persistent protection for a longer period of time.

## Acknowledgments

A portion of cryo-EM data collection was performed at the National Center for Cryo-EM Access and Training and the Simons Electron Microscopy Center located at the New York Structural Biology Center, supported by the NIH Common Fund Transformative High Resolution Cryo-Electron Microscopy program (U24 GM129539) and by grants from the Simons Foundation (SF349247) and NY State Assembly. We thank R. Grassucci, Y.-C. Chi and Z. Zhang from the Cryo-EM Center at Columbia University for assistance with additional cryo-EM data collection. Support for this work was also provided by Samuel Yin, Pony Ma, Peggy & Andrew Cherng, Brii Bioscieces, Jack Ma Foundation, JBP Foundation, Carol Ludwig, and Roger & David Wu, COVID-19 Fast Grants, the Self Graduate Fellowship Program, and NIH grants DP5OD023118, R21AI143407, and R21AI144408.

## Author Contributions

GC and MR determined the cryo-EM structures of 1-57 and 2-7, YG performed bioinformatics analysis. ER and JB produced spike and Fabs. JY, PW, LL, YH, DDH contributed unpublished data related to the neutralization of 1-57 and 2-7 and also produced IgGs and Fabs for these antibodies. DDH supervised IgG production and neutralization assay, PDK contributed to bioinformatics and structural analysis, ZS supervised bioinformatics analysis, LS supervised structural determinations and led the overall project. GC, MR, YG, PDK, ZS and LS wrote the manuscript, with all authors comments.

## Declaration of Interests

DDH, YH, JY, LL and PW are inventors of a patent describing some of the antibodies reported on here.

## RESOURCE AVAILABILITY

### Lead Contact

Further information and requests for resources and reagents should be directed to and will be fulfilled by Lawrence Shapiro.

### Materials Availability

Expression plasmids generated in this study for expressing SARS-CoV-2 proteins and antibodies will be shared upon request.

### Data and Code Availability

The cryo-EM structures have been deposited to the Electron Microscopy Data Bank (EMDB) and the Protein Data Bank (RCSB PDB).

Cryo-EM structural models and maps for antibodies 1-57 and 2-7 in complex with SARS-CoV-2 spike have been deposited in the PDB and EMDB with accession codes PDB 7LS9, EMD-23506, and PDB 7LSS, EMD-23507, respectively.

## EXPERIMENTAL MODEL AND SUBJECT DETAILS

### Cell lines

Expi293F™ Cells (cat# A39240) and Expi293F™ GnTI-Cells (cat# A39240) were from Thermo Fisher Scientific.

### Materials and Methods

#### SARS-CoV-2 spike expression and purification

SARS-CoV-2 S2P spike was produced as described in (Wrapp et al., 2020). Protein expression was carried out in Human Embryonic Kidney (HEK) 293 Freestyle cells (Invitrogen) in suspension culture using serum-free media (Invitrogen) by transient transfection using polyethyleneimine (Polysciences). Cell growths were harvested four days after transfection, and the secreted protein was purified from supernatant by nickel affinity chromatography using Ni-NTA IMAC Sepharose 6 Fast Flow resin (GE Healthcare) followed by size exclusion chromatography on a Superdex 200 column (GE Healthcare) in 10 mM Tris, 150 mM NaCl, pH 7.4.

#### Production of 1-57 and 2-7 Fab

Monoclonal antibody 1-57 was expressed and purified as Fab: VHCH1 with a C-terminal His-tag (His_8_) and LC were constructed separately into the gWiz expression vector, and then co-transfected and expressed in Expi293. Five days after transfection, supernatants were harvested and 1-57 Fab was purified by nickel affinity chromatography using Ni-NTA agarose (Invitrogen cat. No R901-15).

Monoclonal antibody 2-7 was expressed and purified as described in (Liu et al., 2020). Fab fragment was produced by digestion of IgG with immobilized papain at 37 °C for 3 hrs in 50 mM phosphate buffer, 120 mM NaCl, 30 mM cysteine, 1 mM EDTA, pH 7. The resulting Fab was purified from Fc by affinity chromatography on protein A.

Fab purity was analyzed by SDS-PAGE; all Fabs were buffer-exchanged into 10 mM Tris, 150 mM, pH 7.4 for cryo-EM experiments.

#### Cryo-EM samples preparation

The final sample for EM analysis of the 1-57 in complex with SARS-CoV-2 S2P spike was produced by mixing the Fab and spike in a 1:9 molar ratio, with a final trimer concentration of 0.33 mg/mL, followed by incubation on ice for 1 hr. The final buffer was 10 mM sodium acetate, 150 mM NaCl, 0.005% (w/v) n-Dodecyl β-D-maltoside, pH 4.5. Cryo-EM grids were prepared by applying 2 µL of sample to a freshly glow-discharged carbon-coated copper grid (CF 1.2/1.3 300 mesh); the sample was vitrified in liquid ethane using a Vitrobot Mark IV with a wait time of 30 s and a blot time of 3 s.

The final sample for EM analysis of Fab 2-7 in complex with SARS-CoV-2 S2P spike was produced by mixing the Fab and spike in a 1:9 molar ratio, with a final trimer concentration of 0.66 mg/mL, followed by incubation on ice for 1 hr. The final puffer was 10 mM sodium acetate, 150 mM NaCl, 0.005% (w/v) n-Dodecyl β-D-maltoside, pH 5.5. Cryo-EM grids were prepared by applying 2 µL of sample to a freshly plasma-cleaned carbon-coated copper grid (CF 1.2/1.3 300 mesh); the sample was vitrified in liquid ethane using a Leica EMGP with a wait time of 15 s and a blot time of 1.5 s.

#### Cryo-EM data collection, processing and structure refinement

For 1-57, cryo-EM data were collected using the Leginon software (Suloway et al., 2005) installed on a Titan Krios electron microscope operating at 300 kV, equipped with a Gatan K3-BioQuantum direct detection device. The total dose was fractionated for 3 s over 60 raw frames. Motion correction, CTF estimation, particle extraction, 2D classification, ab initio model generation, 3D refinements and local resolution estimation for all datasets were carried out in cryoSPARC 2.15 (Punjani et al., 2017); particles were picked using Topaz (Bepler et al., 2019). The final 3D reconstruction was obtained using non-uniform refinement with C3 symmetry. SARS CoV-2 S2P spike density was modeled using PDB entry 7L2E (Cerutti et al., 2021), as initial template. The initial model for 1-57 Fab variable region was obtained using the SAbPred server (Dunbar et al., 2016).

For 2-7, cryo-EM data were collected as described for 1-57, except that the total electron flux was fractionated over 2 s with 40 total frames. Data processing was also performed as described above. The final reconstruction was obtained using non-uniform refinement with C1 symmetry, followed by local refinement of the ‘down’ RBD and Fab. The SARS-CoV-2 S2P spike density was modeled using PDB entry 6XEY (Liu et al. 2020) as an initial template. A homology model for the 2-7 Fab variable region was obtained using Schrodinger Release 2020-2: BioLuminate (Zhu et al., 2014).

Automated and manual model building were iteratively performed using real space refinement in Phenix (Adams et al., 2004) and Coot (Emsley and Cowtan, 2004) respectively. For 2-7, ISOLDE v1.1 (Croll, 2018) was also used to interactively refine the structure. Half maps were provided to Resolve Cryo-EM tool in Phenix to support manual model building. Geometry validation and structure quality assessment were performed using EMRinger (Barad et al., 2015) and Molprobity (Davis et al., 2004). Map-fitting cross correlation (Fit-in-Map tool) and figures preparation were carried out using PyMOL and UCSF Chimera (Pettersen et al., 2004) and Chimera X (Pettersen et al., 2021). A summary of the cryo-EM data collection, reconstruction and refinement statistics is shown in Table S1.

#### Clustering of published RBD-directed antibodies

Information of published SARS-CoV-2 neutralizing antibodies were obtained from CoV-AbDab database (Raybould et al., 2020), the structure for each antibody was download from PBD. For each pair of antibodies, the RBDs were superimposed and the RMSD of Cα between antibody variable domains were calculated. The heavy and light chain sequence alignment were performed using the online muscle program included in bio3d package (Grant et al., 2006). RMSD of Ca were then calculated based on the sequence alignment by an in-house python script, clustering of the RMSD matrix was performed by hclust package in R.

#### Calculation of antibody targeting frequency for RBD

The epitope RBD-directed antibodies were determined by PISA with the default parameters (Krissinel and Henrick, 2007), the RBD residues with non-zero BSA value were considering as epitope residue. For each residue on RBD, the number of contact antibodies was counted as the frequency of antibody recognition. The antibody targeting frequency was set as b factor of RBD and displayed by Pymol 2.3.2 (DeLano, 2002).

#### Calculation of positional mutation frequency and correlations

The positional mutation frequency were calculated based on the SARS-CoV2 spike sequences deposited in GISAID at Jan 23^rd^, 2021 (Elbe and Buckland-Merrett, 2017). Briefly, spike sequences were aligned pairwise with Wuhan-Hu-01 strain as reference, the mutation frequency for each position were calculated as the total mutations divided by total number of deposited sequences, the low quality ‘X’ residue were not counted as mutations. Normalized effects on ACE2 binding affinity for each mutation on RBD were download from the source data (Starr et al., 2020). The r value and p value for correlations were calculated by cor.test function in R 4.0.3.

## QUANTIFICATION AND STATISTICAL ANALYSIS

The statistical analyses for the pseudovirus neutralization assessments were performed using GraphPad Prism. The SPR data were fitted using Biacore Evaluation Software. Cryo-EM data were processed and analyzed using cryoSPARC. Cryo-EM and crystallographic structural statistics were analyzed using Phenix, Molprobity, EMringer and Chimera. The correlations were performed in R. Statistical details of experiments are described in Method Details or figure legends.

## References

Adams, P.D., Gopal, K., Grosse-Kunstleve, R.W., Hung, L.W., Ioerger, T.R., McCoy, A.J., Moriarty, N.W., Pai, R.K., Read, R.J., Romo, T.D., et al. (2004). Recent developments in the PHENIX software for automated crystallographic structure determination. J Synchrotron Radiat 11, 53–55.

Andrew Rambaut, Nick Loman, Oliver Pybus, Wendy Barclay, Jeff Barrett, Alesandro Carabelli, Tom Connor, Tom Peacock, David L Robertson, and Volz, E. (2020). Preliminary genomic characterisation of an emergent SARS-CoV-2 lineage in the UK defined by a novel set of spike mutations.

Avanzato, V.A., Matson, M.J., Seifert, S.N., Pryce, R., Williamson, B.N., Anzick, S.L., Barbian, K., Judson, S.D., Fischer, E.R., Martens, C., et al. (2020). Case Study: Prolonged Infectious SARS-CoV-2 Shedding from an Asymptomatic Immunocompromised Individual with Cancer. Cell 183, 1901–1912 e1909.

Banach, B.B., Cerutti, G., Fahad, A.S., Shen, C.-H., de Souza, M.O., Katsamba, P.S., Tsybovsky, Y., Wang, P., Nair, M.S., Huang, Y., et al. (2021). Paired heavy and light chain signatures contribute to potent SARS-CoV-2 neutralization in public antibody responses. bioRxiv, 2020.2012.2031.424987.

Barad, B.A., Echols, N., Wang, R.Y., Cheng, Y., DiMaio, F., Adams, P.D., and Fraser, J.S. (2015). EMRinger: side chain-directed model and map validation for 3D cryo-electron microscopy. Nat Methods 12, 943–946.

Baric, R.S. (2020). Emergence of a Highly Fit SARS-CoV-2 Variant. N Engl J Med 383, 2684–2686.

Barnes, C.O., Jette, C.A., Abernathy, M.E., Dam, K.A., Esswein, S.R., Gristick, H.B., Malyutin, A.G., Sharaf, N.G., Huey-Tubman, K.E., Lee, Y.E., et al. (2020). SARS-CoV-2 neutralizing antibody structures inform therapeutic strategies. Nature 588, 682–687.

Baum, A., Fulton, B.O., Wloga, E., Copin, R., Pascal, K.E., Russo, V., Giordano, S., Lanza, K., Negron, N., Ni, M., et al. (2020). Antibody cocktail to SARS-CoV-2 spike protein prevents rapid mutational escape seen with individual antibodies. Science 369, 1014–1018.

Bepler, T., Morin, A., Rapp, M., Brasch, J., Shapiro, L., Noble, A.J., and Berger, B. (2019). Positive-unlabeled convolutional neural networks for particle picking in cryo-electron micrographs. Nat Methods 16, 1153–1160.

Brouwer, P.J.M., Caniels, T.G., van der Straten, K., Snitselaar, J.L., Aldon, Y., Bangaru, S., Torres, J.L., Okba, N.M.A., Claireaux, M., Kerster, G., et al. (2020). Potent neutralizing antibodies from COVID-19 patients define multiple targets of vulnerability. Science 369, 643–650.

Callaway, E., Cyranoski, D., Mallapaty, S., Stoye, E., and Tollefson, J. (2020). The coronavirus pandemic in five powerful charts. Nature 579, 482–483.

Cerutti, G., Guo, Y., Zhou, T., Gorman, J., Lee, M., Rapp, M., Reddem, E.R., Yu, J., Bahna, F., Bimela, J., et al. (2021). Potent SARS-CoV-2 Neutralizing Antibodies Directed Against Spike N-Terminal Domain Target a Single Supersite. bioRxiv, 2021.2001.2010.426120.

Chi, X., Yan, R., Zhang, J., Zhang, G., Zhang, Y., Hao, M., Zhang, Z., Fan, P., Dong, Y., Yang, Y., et al. (2020). A neutralizing human antibody binds to the N-terminal domain of the Spike protein of SARS-CoV-2. Science, eabc6952.

Choi, B., Choudhary, M.C., Regan, J., Sparks, J.A., Padera, R.F., Qiu, X., Solomon, I.H., Kuo, H.H., Boucau, J., Bowman, K., et al. (2020). Persistence and Evolution of SARS-CoV-2 in an Immunocompromised Host. N Engl J Med 383, 2291–2293.

Croll, T.I. (2018). ISOLDE: a physically realistic environment for model building into low-resolution electron-density maps. Acta Crystallogr D Struct Biol 74, 519–530.

Cucinotta, D., and Vanelli, M. (2020). WHO Declares COVID-19 a Pandemic. Acta Biomed 91, 157–160.

Davis, I.W., Murray, L.W., Richardson, J.S., and Richardson, D.C. (2004). MOLPROBITY: structure validation and all-atom contact analysis for nucleic acids and their complexes. Nucleic Acids Res 32, W615–619.

DeLano, W.L. (2002). The PyMOL Molecular Graphics System (San Carlos, CA: DeLano Scientific).

Dong, E., Du, H., and Gardner, L. (2020). An interactive web-based dashboard to track COVID-19 in real time. Lancet Infect Dis 20, 533–534.

Dunbar, J., Krawczyk, K., Leem, J., Marks, C., Nowak, J., Regep, C., Georges, G., Kelm, S., Popovic, B., and Deane, C.M. (2016). SAbPred: a structure-based antibody prediction server. Nucleic Acids Res 44, W474–478.

Elbe, S., and Buckland-Merrett, G. (2017). Data, disease and diplomacy: GISAID’s innovative contribution to global health. Glob Chall 1, 33–46.

Emsley, P., and Cowtan, K. (2004). Coot: model-building tools for molecular graphics. Acta Crystallogr D Biol Crystallogr 60, 2126–2132.

Grant, B.J., Rodrigues, A.P., ElSawy, K.M., McCammon, J.A., and Caves, L.S. (2006). Bio3d: an R package for the comparative analysis of protein structures. Bioinformatics 22, 2695-2696.

Hansen, J., Baum, A., Pascal, K.E., Russo, V., Giordano, S., Wloga, E., Fulton, B.O., Yan, Y., Koon, K., Patel, K., et al. (2020). Studies in humanized mice and convalescent humans yield a SARS-CoV-2 antibody cocktail. Science, 10.1126/science.abd0827.

Krissinel, E., and Henrick, K. (2007). Inference of macromolecular assemblies from crystalline state. J Mol Biol 372, 774–797.

Ku, Z., Xie, X., Davidson, E., Ye, X., Su, H., Menachery, V.D., Li, Y., Yuan, Z., Zhang, X., Muruato, A.E., et al. (2021). Molecular determinants and mechanism for antibody cocktail preventing SARS-CoV-2 escape. Nat Commun 12, 469.

Liu, L., Wang, P., Nair, M.S., Yu, J., Rapp, M., Wang, Q., Luo, Y., Chan, J.F., Sahi, V., Figueroa, A., et al. (2020). Potent neutralizing antibodies against multiple epitopes on SARS-CoV-2 spike. Nature 584, 450–456.

Lv, Z., Deng, Y.Q., Ye, Q., Cao, L., Sun, C.Y., Fan, C., Huang, W., Sun, S., Sun, Y., Zhu, L., et al. (2020). Structural basis for neutralization of SARS-CoV-2 and SARS-CoV by a potent therapeutic antibody. Science 369, 1505–1509.

Nuno R. Faria, Ingra Morales Claro, Darlan Candido, Lucas A. Moyses Franco, Pamela S. Andrade, Thais M. Coletti, Camila A. M. Silva, Flavia C. Sales, Erika R. Manuli, Renato S. Aguiar, et al. (2021). Genomic characterisation of an emergent SARS-CoV-2 lineage in Manaus: preliminary findings.

Pettersen, E.F., Goddard, T.D., Huang, C.C., Couch, G.S., Greenblatt, D.M., Meng, E.C., and Ferrin, T.E. (2004). UCSF Chimera--a visualization system for exploratory research and analysis. J Comput Chem 25, 1605–1612.

Pettersen, E.F., Goddard, T.D., Huang, C.C., Meng, E.C., Couch, G.S., Croll, T.I., Morris, J.H., and Ferrin, T.E. (2021). UCSF ChimeraX: Structure visualization for researchers, educators, and developers. Protein Sci 30, 70–82.

Pinto, D., Park, Y.J., Beltramello, M., Walls, A.C., Tortorici, M.A., Bianchi, S., Jaconi, S., Culap, K., Zatta, F., De Marco, A., et al. (2020). Cross-neutralization of SARS-CoV-2 by a human monoclonal SARS-CoV antibody. Nature 583, 290–295.

Punjani, A., Rubinstein, J.L., Fleet, D.J., and Brubaker, M.A. (2017). cryoSPARC: algorithms for rapid unsupervised cryo-EM structure determination. Nat Methods 14, 290–296.

Rapp, M., Guo, Y., Reddem, E.R., Liu, L., Wang, P., Yu, J., Cerutti, G., Bimela, J., Bahna, F., Mannepalli, S., et al. (2021). Modular basis for potent SARS-CoV-2 neutralization by a prevalent VH1-2-derived antibody class. bioRxiv, 2021.2001.2011.426218.

Raybould, M.I.J., Kovaltsuk, A., Marks, C., and Deane, C.M. (2020). CoV-AbDab: the Coronavirus Antibody Database. Bioinformatics.

Robbiani, D.F., Gaebler, C., Muecksch, F., Lorenzi, J.C.C., Wang, Z., Cho, A., Agudelo, M., Barnes, C.O., Gazumyan, A., Finkin, S., et al. (2020). Convergent antibody responses to SARS-CoV-2 in convalescent individuals. Nature 584, 437–442.

Sabino, E.C., Buss, L.F., Carvalho, M.P.S., Prete, C.A., Jr., Crispim, M.A.E., Fraiji, N.A., Pereira, R.H.M., Parag, K.V., da Silva Peixoto, P., Kraemer, M.U.G., et al. (2021). Resurgence of COVID-19 in Manaus, Brazil, despite high seroprevalence. Lancet 397, 452–455.

Shi, R., Shan, C., Duan, X., Chen, Z., Liu, P., Song, J., Song, T., Bi, X., Han, C., Wu, L., et al. (2020). A human neutralizing antibody targets the receptor-binding site of SARS-CoV-2. Nature, 10.1038/s41586-41020-42381-y.

Shrock, E., Fujimura, E., Kula, T., Timms, R.T., Lee, I.H., Leng, Y., Robinson, M.L., Sie, B.M., Li, M.Z., Chen, Y., et al. (2020). Viral epitope profiling of COVID-19 patients reveals cross-reactivity and correlates of severity. Science 370.

Starr, T.N., Greaney, A.J., Hilton, S.K., Ellis, D., Crawford, K.H.D., Dingens, A.S., Navarro, M.J., Bowen, J.E., Tortorici, M.A., Walls, A.C., et al. (2020). Deep Mutational Scanning of SARS-CoV-2 Receptor Binding Domain Reveals Constraints on Folding and ACE2 Binding. Cell 182, 1295–1310 e1220.

Suloway, C., Pulokas, J., Fellmann, D., Cheng, A., Guerra, F., Quispe, J., Stagg, S., Potter, C.S., and Carragher, B. (2005). Automated molecular microscopy: the new Leginon system. J Struct Biol 151, 41–60.

Tegally, H., Wilkinson, E., Giovanetti, M., Iranzadeh, A., Fonseca, V., Giandhari, J., Doolabh, D., Pillay, S., San, E.J., Msomi, N., et al. (2020). Emergence and rapid spread of a new severe acute respiratory syndrome-related coronavirus 2 (SARS-CoV-2) lineage with multiple spike mutations in South Africa. medRxiv, 2020.2012.2021.20248640.

Tortorici, M.A., Beltramello, M., Lempp, F.A., Pinto, D., Dang, H.V., Rosen, L.E., McCallum, M., Bowen, J., Minola, A., Jaconi, S., et al. (2020). Ultrapotent human antibodies protect against SARS-CoV-2 challenge via multiple mechanisms. Science 370, 950–957.

Wang, P., Liu, L., Iketani, S., Luo, Y., Guo, Y., Wang, M., Yu, J., Zhang, B., Kwong, P.D., Graham, B.S., et al. (2021). Increased Resistance of SARS-CoV-2 Variants B.1.351 and B.1.1.7 to Antibody Neutralization. bioRxiv, 2021.2001.2025.428137.

West, A.P., Barnes, C.O., Yang, Z., and Bjorkman, P.J. (2021). SARS-CoV-2 lineage B.1.526 emerging in the New York region detected by software utility created to query the spike mutational landscape. bioRxiv, 2021.2002.2014.431043.

Wibmer, C.K., Ayres, F., Hermanus, T., Madzivhandila, M., Kgagudi, P., Lambson, B.E., Vermeulen, M., van den Berg, K., Rossouw, T., Boswell, M., et al. (2021). SARS-CoV-2 501Y.V2 escapes neutralization by South African COVID-19 donor plasma. bioRxiv, 021.2001.2018.427166.

Wrapp, D., Wang, N., Corbett, K.S., Goldsmith, J.A., Hsieh, C.L., Abiona, O., Graham, B.S., and McLellan, J.S. (2020). Cryo-EM structure of the 2019-nCoV spike in the prefusion conformation. Science 367, 1260–1263.

Wu, Y., Wang, F., Shen, C., Peng, W., Li, D., Zhao, C., Li, Z., Li, S., Bi, Y., Yang, Y., et al. (2020). A noncompeting pair of human neutralizing antibodies block COVID-19 virus binding to its receptor ACE2. Science 368, 1274–1278.

Yuan, M., Liu, H., Wu, N.C., Lee, C.D., Zhu, X., Zhao, F., Huang, D., Yu, W., Hua, Y., Tien, H., et al. (2020). Structural basis of a shared antibody response to SARS-CoV-2. Science 369, 1119–1123.

Zhu, K., Day, T., Warshaviak, D., Murrett, C., Friesner, R., and Pearlman, D. (2014). Antibody structure determination using a combination of homology modeling, energy-based refinement, and loop prediction. Proteins 82, 1646–1655.

