## Supplemental Figures for "Structural Basis for Accommodation of Emerging B.1.351 and B.1.1.7 Variants by Two Potent SARS-CoV-2 Neutralizing Antibodies"

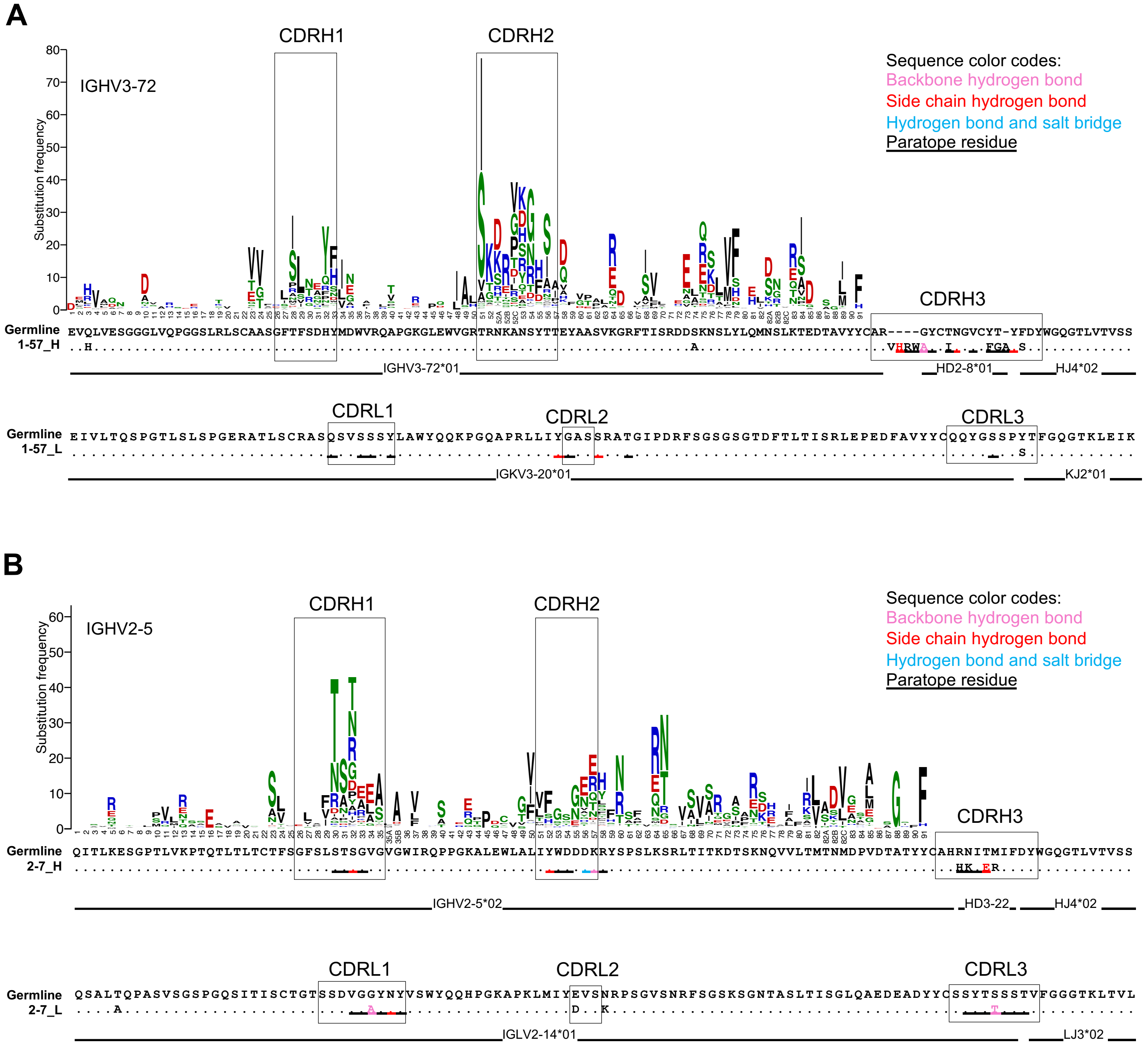


**Figure S1. Sequence alignment for 1-57 and 2-7 with their corresponding germline genes, Related to Figures 1- 2**

(A) 1-57 heavy and light chain aligned with VH3-72*01 and IGKV3-20*01, the gene-specific substitution profile (GSSP) showing somatic hypermutation probabilities for VH3-72 gene.

(B) 2-7 heavy and light chain aligned with VH2-5*02 and IGLV2-14*01, the gene-specific substitution profile (GSSP) showing somatic hypermutation probabilities for VH2-5 gene.

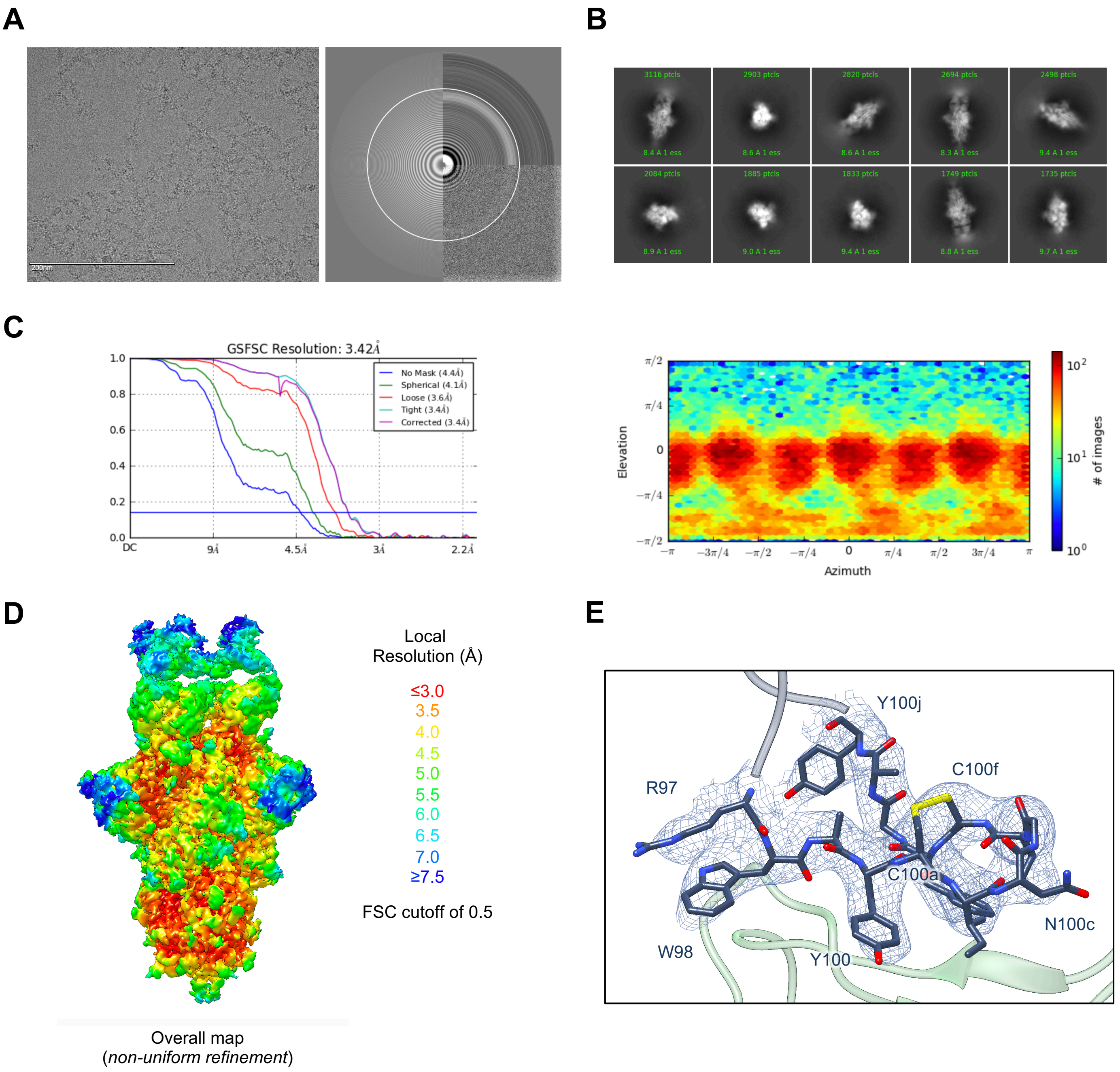


**Figure S2. Cryo-EM details of 1-57 Fab in complex with SARS-CoV-2 S2P spike, Related to Figure 1.**
(A) Representative micrograph and CTF of the micrograph are shown.
(B) Representative 2D class averages are shown.
(C) The gold-standard Fourier shell correlation resulted in a resolution of 3.42 Å for the overall map using non-uniform refinement with C3 symmetry (left panel); the orientations of all particles used in the final refinement are shown as a heatmap (right panel).
(D) The local resolution of the final overall map and locally refined map are shown, generated through cryoSPARC using an FSC cutoff of 0.5.
(E) Representative density is shown for the CDR H3 loop of 1-57 contacting RBD; the contour level is 0.25 (1.6σ). CDR H3 carbon atoms are colored in dark blue, oxygen in red, nitrogen in blue; RBD is colored in green.

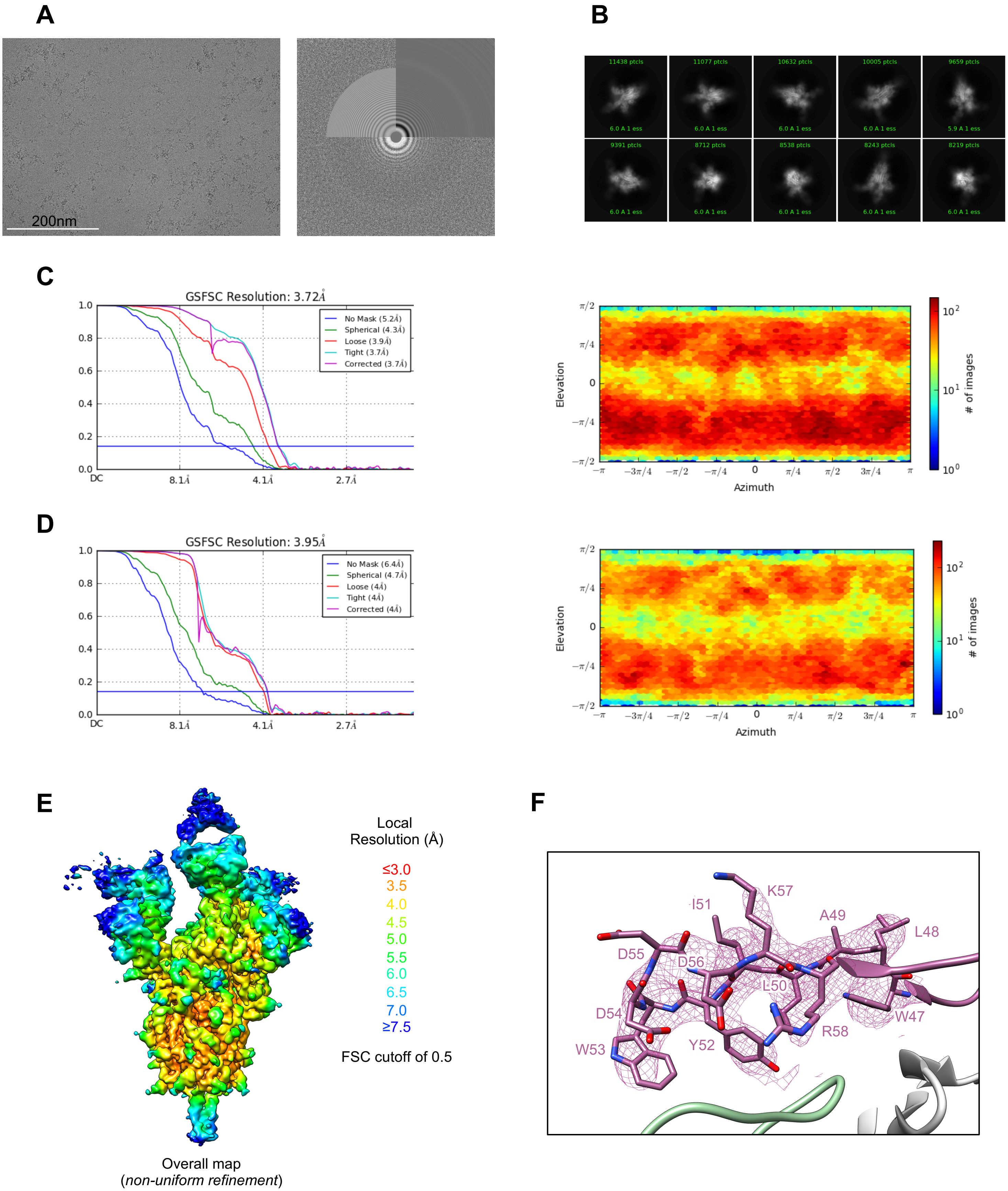


**Figure S3. Cryo-EM details of 2-7 Fab in complex with SARS-CoV-2 S2P spike, Related to Figure 2.**
(A) Representative micrograph and CTF of the micrograph are shown.
(B) Representative 2D class averages are shown.
(C) The gold-standard Fourier shell correlation resulted in a resolution of 3.72 Å for the overall map using non-uniform refinement with C1 symmetry (left panel); the orientations of all particles used in the final refinement are shown as a heatmap (right panel).
(D) The gold-standard Fourier shell correlation resulted in a resolution of 3.95 Å for Fab/RBD interface using local refinement. The orientations of all particles used in the final refinement are shown as a heatmap (right panel).
(E) The local resolution of the final overall map and locally refined map are shown, generated through cryoSPARC using an FSC cutoff of 0.5.
(F) Representative density is shown for the CDR H2 loop of 2-7 contacting RBD; the contour level is 0.409 (1.2σ) . CDR H2 carbon atoms are colored in dark purple, oxygen in red, nitrogen in blue; RBD is colored in green, and the light chain in light gray.
