## Supplemental Table for "Structural Basis for Accommodation of Emerging B.1.351 and B.1.1.7 Variants by Two Potent SARS-CoV-2 Neutralizing Antibodies"

**Table S1. Cryo-EM Data Collection and Refinement Statistics, Related to Figures 1-2.**

| SARS-CoV-2 S2P complex | 1-57 Fab | 2-7 Fab |
|---|---|---|
| **EMDB ID** | EMD-23506 | EMD-23507 |
| **PDB ID** | 7LS9 | 7LSS |
| Data Collection | | |
| Microscope | FEI Titan Krios | FEI Titan Krios |
| Voltage (kV) | 300 | 300 |
| Electron dose ($e^-/Å^2$) | 41.92 | 51.69 |
| Detector | Gatan K3 BioQuantum | Gatan K3 BioQuantum |
| Pixel Size (Å) | 1.07 | 1.058 |
| Defocus Range (μm) | -0.8/-2.5 | -0.5/-2.5 |
| Magnification | 81000 | 81000 |
| Reconstruction | | |
| Software | cryoSPARC v2.15 | cryoSPARC v2.15 |
| Particles | 89,601 | 165,576 |
| Symmetry | C3 | C1 |
| Box size (pix) | 420 | 384 |
| Resolution (Å) ($FSC_{0.143}$) | 3.42 | 3.72 |
| Refinement | | |
| Software | Phenix 1.18 | Phenix 1.18 |
| Protein residues | 4002 | 2783 |
| Chimera CC | 0.83 | 0.79 |
| EMRinger Score | 2.14 | 3.13 |
| R.m.s. deviations | | |
| Bond lengths (Å) | 0.006 | 0.011 |
| Bond angles (°) | 1.11 | 1.066 |
| Validation | | |
| Molprobity score | 1.38 | 1.15 |
| Clash score | 4.32 | 1.08 |
| Favored rotamers (%) | 100 | 100 |
| Ramachandran | | |
| Favored regions (%) | 97.0 | 95.09 |
| Allowed regions (%) | 3.0 | 4.91 |
| Disallowed regions (%) | 0 | 0 |
